# Real-world size of objects serves as an axis of object space

**DOI:** 10.1101/2021.09.28.462153

**Authors:** Taicheng Huang, Yiying Song, Jia Liu

## Abstract

Our mind can represent various objects from the physical world metaphorically into an abstract and complex high-dimensional object space, with a finite number of orthogonal axes encoding critical object features. Previous fMRI studies have shown that the middle fusiform sulcus in the ventral temporal cortex separates the real-world small-size map from the large-size map. Here we asked whether the feature of objects’ real-world size constructed an axis of object space with deep convolutional neural networks (DCNNs) based on three criteria of sensitivity, independence and necessity that are impractical to be examined altogether with traditional approaches. A principal component analysis on features extracted by the DCNNs showed that objects’ real-world size was encoded by an independent component, and the removal of this component significantly impaired DCNN’s performance in recognizing objects. By manipulating stimuli, we found that the shape and texture of objects, rather than retina size, co-occurrence and task demands, accounted for the representation of the real-world size in the DCNNs. A follow-up fMRI experiment on humans further demonstrated that the shape, but not the texture, was used to infer the real-world size of objects in humans. In short, with both computational modeling and empirical human experiments, our study provided the first evidence supporting the feature of objects’ real-world size as an axis of object space, and devised a novel paradigm for future exploring the structure of object space.

**Teaser:** This work provides the first evidence illuminating the feature of objects’ real-world size as an axis of the object space for object recognition with a mutually-inspired paradigm of computational modelling and biological observation.

## Introduction

Objects are complicated, and humans have evolved an excellent ability to recognize them in complex environments. One possible underlying mechanism for such feat is to extract object features to form an object space, where each axis carries critical and orthogonal information regarding one aspect of object properties (*1–3*). Thus, object recognition is considered as a computational problem of finding a division surface to separate different objects represented by features projected from axes (*4, 5*). This hypothetical object space is likely implemented in primate’s inferotemporal cortex, as two orthogonal axes, the animacy (animate versus inanimate) and curvature (spiky versus stubby), have been identified, which classify macaque’s ventral temporal cortex into four distinct clusters based on neuronal responses to these two features (*6*). However, these two features are not sufficient to represent all object categories, as tennis balls and the earth are both inanimate and stubby, but they belong to two object categories. Here, in this study we tried to identify more features that serve as axes of object space.

One potential candidate is the real-world size of objects that provides critical information for embodied cognition. First, the real-world size described the scale of an object in the natural environment, which implies the potential layout of objects (*7*). For example, the size of whales suggests its co-occurrence with seas but not creeks. Second, the real-world size of objects provides heuristic information for affordance, as big objects are less likely manipulatable (*8*). Third, fMRI studies on humans have revealed that the real-world size of objects is encoded in human’s ventral temporal cortex, where the medial part of the occipitotemporal cortex is preferable to object with a larger size and the lateral to a smaller size (*9*). Finally, size information is apparently encoded independently from animacy, and they together form a tripartite organizational schema (i.e., big objects, all animals and small objects) in the ventral occipitotemporal cortex (*10*) and medial temporal lobe (*11*).

However, the neuronal sensitivity (sensitivity criterion) to a feature is requisite, but not sufficient, for examining whether the feature serves as an axis of object space. At least two more criteria need to be satisfied. First, the feature shall be statistically orthogonal to other features (independence criterion). Second, the removal of this feature will significantly impair the performance of object recognition (necessity criterion). Unfortunately, previous studies that tried to identify axes of object space do not directly test these two criteria, simply because it is impractical with traditional methods on monkeys and human subjects. Indeed, the criterion of statistical independence requires recording of neuronal responses to a gigantic number of objects so that the independence can be tested between the axis-to-be-identified and the rest of all axes of object space. The criterion of necessity requires the removal of the axis of object space, not the image space, to examine the behavioral performance based on the deficit representation. This is virtually impractical in neurophysiological or fMRI studies.

Recently, deep convolutional neural networks (DCNNs) show great potentials of simulating primates’ ventral visual pathway on object recognition (*12–16*), with similar representations between DCNNs and primates been revealed (e.g., retinotopy (*17*), semantic structure (*18*), coding scheme (*19*), and face representation (*20*)). More importantly, in DCNNs the object space can be constructed with millions of natural images and flexibly manipulated, which are perfect for testing the criteria of independence and necessity, along with the criterion of sensitivity. Therefore, here we used DCNNs to investigate whether the feature of object’s real-world size served as an axis of object space.

To do this, we first examined whether a typical DCNN, AlexNet (*21*), encoded object size (sensitivity criterion) in an independent fashion (independence criterion), and the removal of such axis impaired the performance of the DCNN (necessity criterion). Second, we systematically manipulated stimuli and explored factors (e.g., the retinal size, co-occurrence, task demands, shape and texture of objects) that may affect DCNN’s extraction of objects’ real-world size. Finally, we used fMRI to examine whether the factors identified in the DCNN also contributed to humans’ representation of objects’ real-world size.

## Results

We first evaluated whether an AlexNet pre-trained with object recognition extracted the feature of objects’ real-world size (sensitivity criterion) by examining the correspondence of AlexNet’s responses and objects’ real-world size. We found that the representational similarity matrix (RSM) of objects with different real-world sizes (Fig. 1A) at the Conv4 layer of AlexNet was highly correlated with the RSM of an ideal observer based on objects’ real-world size (Fig. 1B) (r = 0.96, p < 0.001) and the RSM based on humans’ subjective judgment on objects’ real-world size (Fig. 1C; r = 0.94, p < 0.001), suggesting that the AlexNet extracted the feature of objects’ real-world size in the same way as humans did. The similarity between the RSMs was observed in all convolution layers except the Conv1 layer (Fig. S1). Further investigation found the sensitivity to real-world size was also observed in DCNNs with different architectures (Fig. S2), but disappeared in an untrained AlexNet (see Fig. S3), suggesting that the representation for objects’ real-world size was formed through DCNN training, regardless of physical implementations.

**Fig. 1.**
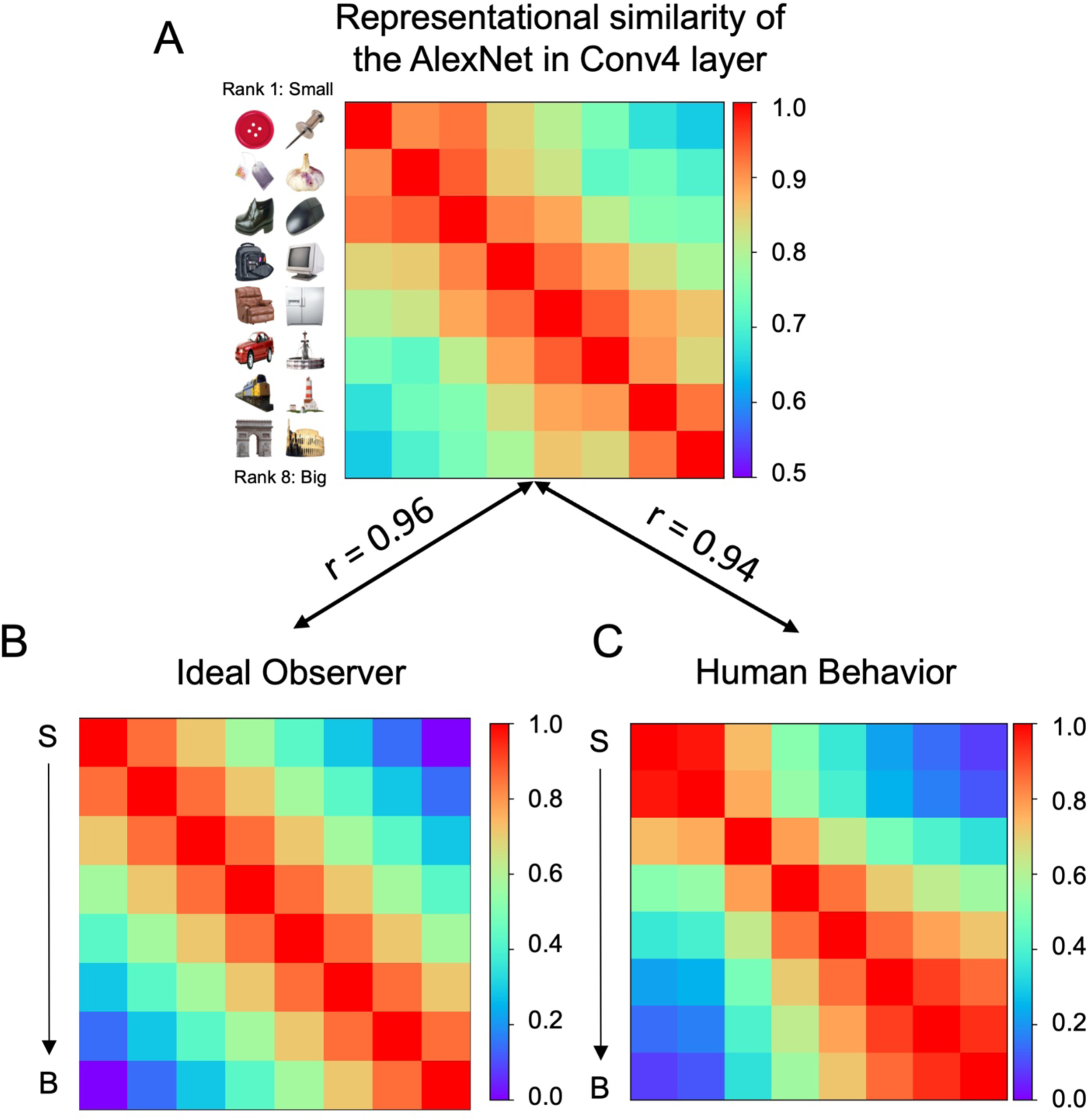
Real-world size preference automatically emerged in the AlexNet designated for object recognition. (A) The representational similarity matrix (RSM) of Conv4’s responses to objects with different sizes. As shown in the Fig., objects with similar real-world sizes elicited similar responses in Conv4. (B) The RSM of the ideal observer based on objects’ real-world size, and (C) the RSM of humans based on subjective judgment. S: small objects. B: big objects.

Since the feature of objects’ real-world was extracted by the AlexNet, we then examined whether it served as an axis of object space that is orthogonal to features encoded by other axes (independence criterion). To do this, we first used principal component analysis (PCA) on AlexNet Conv4’s responses to 50,000 ImageNet validation images to construct an object space with 50 orthogonal axes. By iteratively removing Conv4’s response variance aligned to one axis, we examined the similarity between the RSM of the residual variance and that of the ideal observer. We found that with the removal of the second PC (PC2), the similarity was significantly reduced (Fig. 2A, left), whereas the removal of the rest PCs did not significantly decrease the similarity (Fig. 2A, right. PC2: DI = 0.40, p < 0.05, Bonferroni corrected; the rest: DIs < 0.01, ps > 0.05). This pattern was replicated in DCNNs with different architectures (Fig. S4A). In short, the feature of objects’ real-world size was specifically encoded by one axis of object space alone.

**Fig. 2.**
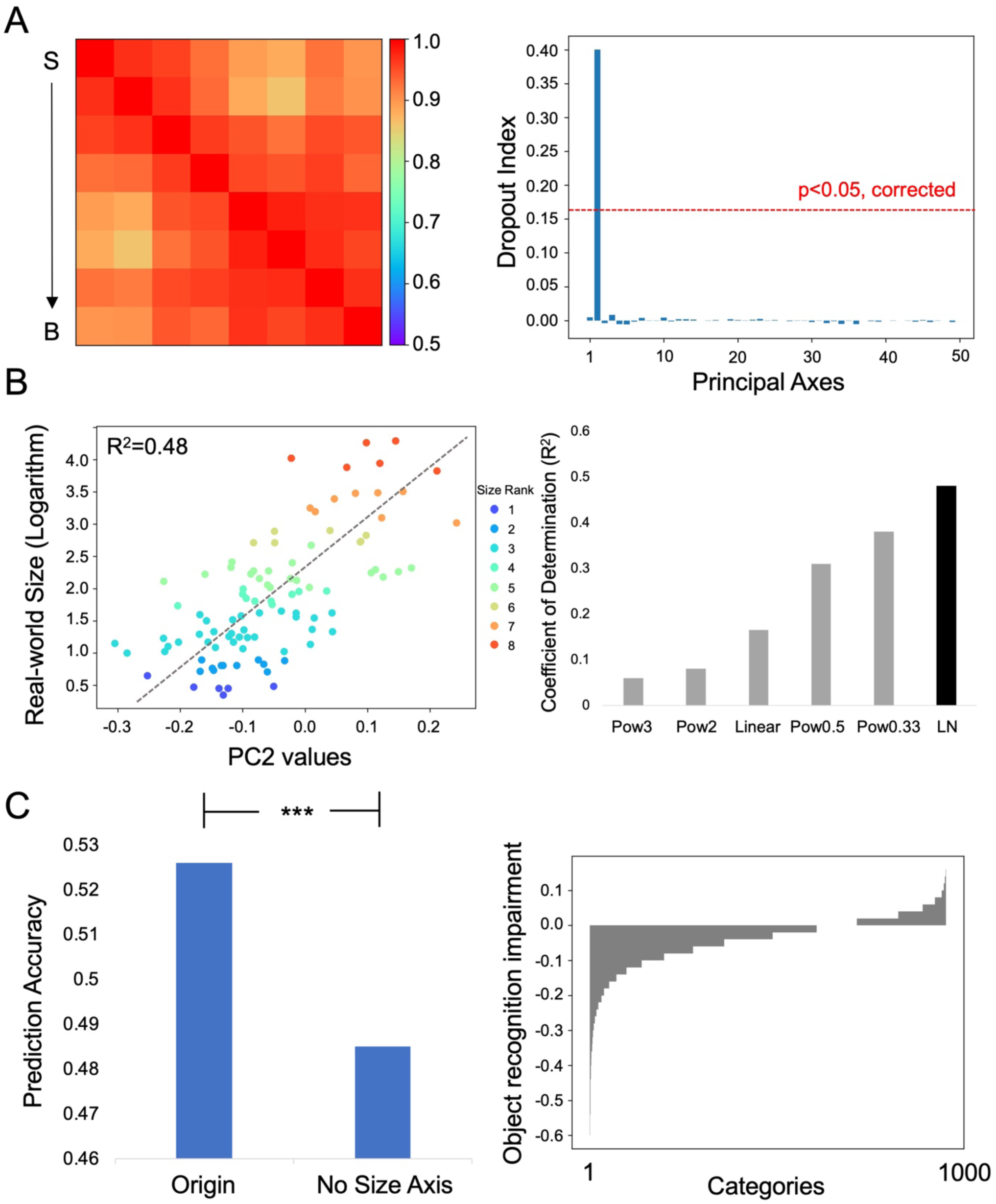
The axis of object space that encoded objects’ real-world size. (A) When the variance that aligned to the second axis was removed from Conv4’s responses, the RSM of Conv4’s responses did not show sensitivity to objects’ real-world size anymore, as compared to that of the ideal observer (left). Specifically, only the removal of the second axis showed a significant decrease in similarity, whereas the rest axes were insensitive to this feature (right). (B) Quantitative mapping of objects’ real-world size to the PC2 was best described with a natural logarithm function (left), among all functions tested (right). The color indicates size rank. (C) The removal of the response variance aligned to the second axis significantly impaired AlexNet’s performance on object recognition in general (left). Classification accuracy of most categories was impaired in varying degrees after the removal of the PC2 (right). S: small objects; B: big objects. Pow: power; LN: natural logarithms.

How did this axis quantitatively encode real-world size? Among all functions tested (e.g., linear, exponential, and logarithm), the best function that map the physical world (i.e., real-world size) to the representational space (i.e., values of PC2) is the natural logarithm scale (R^2^ = 0.48, p < 0.001, Fig. 2B, left), among all functions tested (Fig. 2B, right). Thus, the stimulus-representation mapping apparently followed the Weber-Fechner law, in line with the mapping rule found in humans (*22*). Similar results were also found with other DCNNs (Fig. S4B).

To examine the third criterion of necessity, we tested whether the removal of response variance aligned to this axis significantly impaired AlexNet’s performance on object recognition. To do this, the PC2 was regressed out from Conv4’s responses to the ImageNet validation images, and the residue responses were then fed into further layers. We found that the Top-1 accuracy was significantly decreased from 52.6% to 48.5% (p < 0.001) in general (Fig. 2C, left). A careful examination of the recognition accuracy of each object category showed that the Top-1 accuracies of 637 categories were decreased, 113 categories unchanged and 250 categories increased. Similar results were also found in other DCNNs (Fig. S4C). Taken together, the feature of objects’ real-world size satisfied the three criteria of sensitivity, independence, and necessity to serve as an axis of object space.

After identifying the feature of objects’ real-world size as an axis of object space, a more interesting question is how the DCNN acquired such information. An intuition is that in natural environments, an object is seldomly present alone, thus the relative difference in retinal size among objects in images (e.g., the person in Fig. 3A is smaller than the car) may provide information about objects’ real-world size. To examine this possibility, we trained an AlexNet with images containing only single object without any background (i.e., the single-object AlexNet). Surprisingly, the single-object AlexNet was still able to represent objects’ real-world size, suggested by the close correspondence of Conv4’s responses to the ideal observer (Fig. 3B; sensitivity: r = 0.96, p < 0.001) and by an axis specifically encoding this feature in object space (Fig. 3C, independence: DI=0.30, p<0.05, Bonferroni correction) with natural logarithm function (R^2^=0.37, p<0.001). We also manipulated the retinal size of the objects from the real-world size dataset, and the representation of object’s real-world size remained unchanged (Fig. S5). In short, the encoding of objects’ real-world size unlikely resulted from the relative retinal size differences among objects or the absolute retinal size of objects in images.

**Fig. 3.**
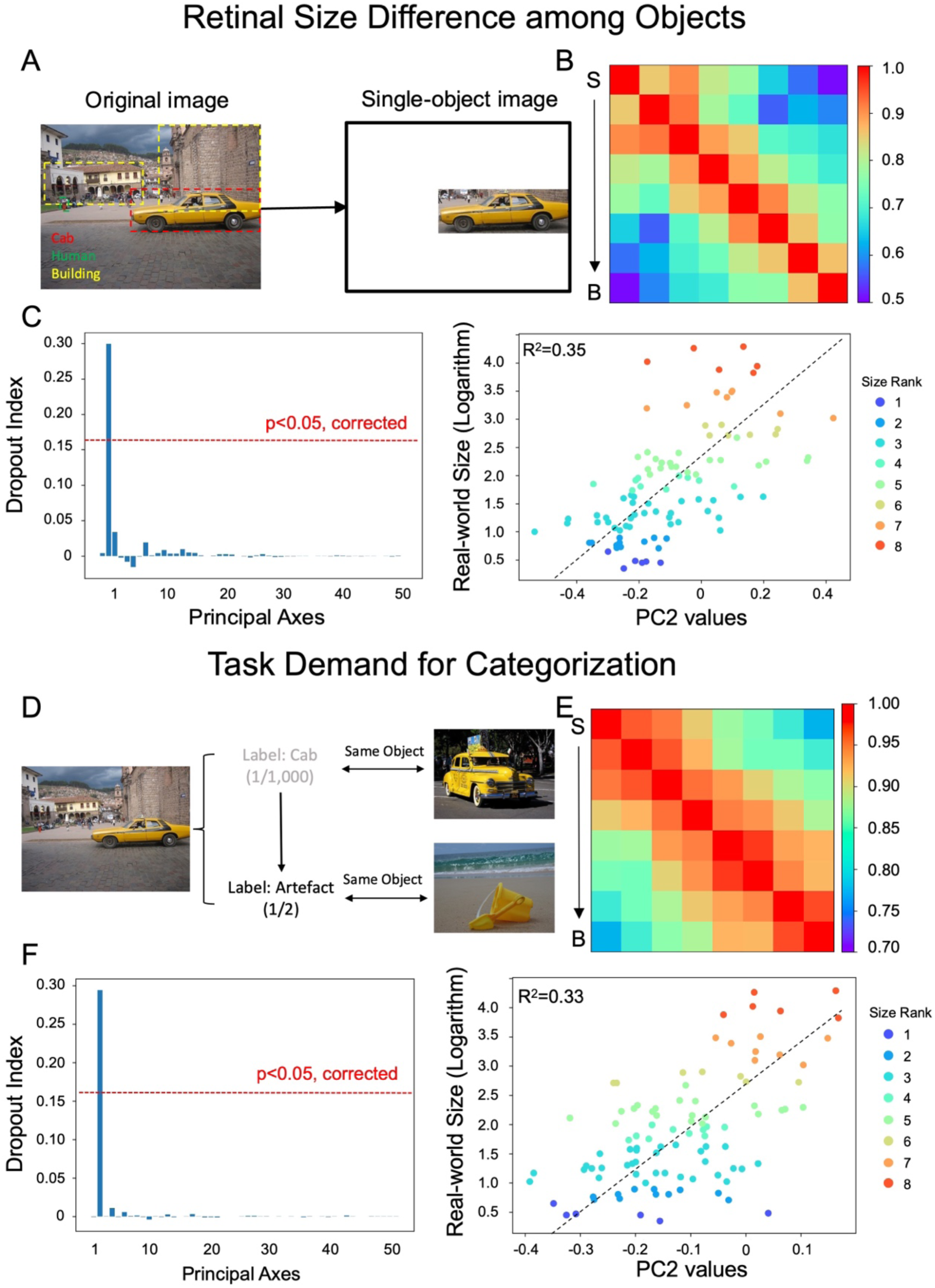
The factors of objects’ retinal size and task demands in representing objects’ real-world size. (A) Differences in retinal size among objects. In natural images, objects with smaller retinal size (e.g., person) are likely smaller than those with larger retinal size (e.g., car, building). The effect of this factor was tested by removing the background. (B) The RSM of Conv4’s responses in the single-object AlexNet to objects’ real-world size. (C) The second axis in object space specifically encoded the real-world size, and the best mapping function was the natural logarithm. (D) Task demands. The task of the AlexNet (i.e., the AlexNet-Cate2) was designated to categorize all objects into two superordinate categories: animate versus artefacts, where buckets were not differentiated from cars, for example. (E) The RSM of Conv4’s responses in the AlexNet-Cate2. (F) The second axis in object space also specifically encoded the real-world size with the mapping function of the natural logarithm. S: small objects; B: big objects.

Another factor is the task demands of the DCNNs. Recent studies have shown that task demands modulate the intrinsic representation of the DCNNs (*18, 20*). To test whether task demands affected the representation of objects’ real-world size, we trained an AlexNet with the same image datasets but to differentiate objects at a superordinate level: animate versus artefacts (i.e., the AlexNet-Cate2). Again, the task demand had little effect on the representation of objects’ real-world size, with the best correspondence to the ideal observer in Conv4 (Fig. 3F; r = 0.95, p < 0.001) and the same axis encoding real-world size of objects (DI = 0.27, p < 0.05, Bonferroni correction) with natural logarithm function (R^2^ = 0.37, p < 0.001) (Fig. 3F).

Since the external factors had little effect on the DCNN to extract the feature of objects’ real-world size to construct object space, here we tested whether the intrinsic factors of objects, shape and texture, were critical. Three types of stimuli were created from the real-world size dataset: silhouette, texture and shuffle images (Fig. 4A). The silhouettes reserve the overall outline, the textures contain local features, and the shuffle images remove both shape and texture information. We found that the AlexNet was able to infer objects’ real-world size with either shape (r = 0.89, p < 0.001) or texture information (r = 0.73, p < 0.001) of objects (Fig. 4B), which was also encoded in the second axis of object space (silhouettes: R^2^ = 0.34, p < 0.001; textures: R^2^ = 0.33, p < 0.001; Fig. 4C). In contrast, when the shape and texture information was scrambled, the AlexNet was no longer able to extract the feature of objects’ real-world size (R^2^ = 0.01, p > 0.05). Therefore, the DCNNs seemed to use the intrinsic properties of objects (i.e., shape and texture) but not the external factors (i.e., co-occurrence, absolute retinal size, and task demands) to represent objects’ real-world size in object space.

**Fig. 4.**
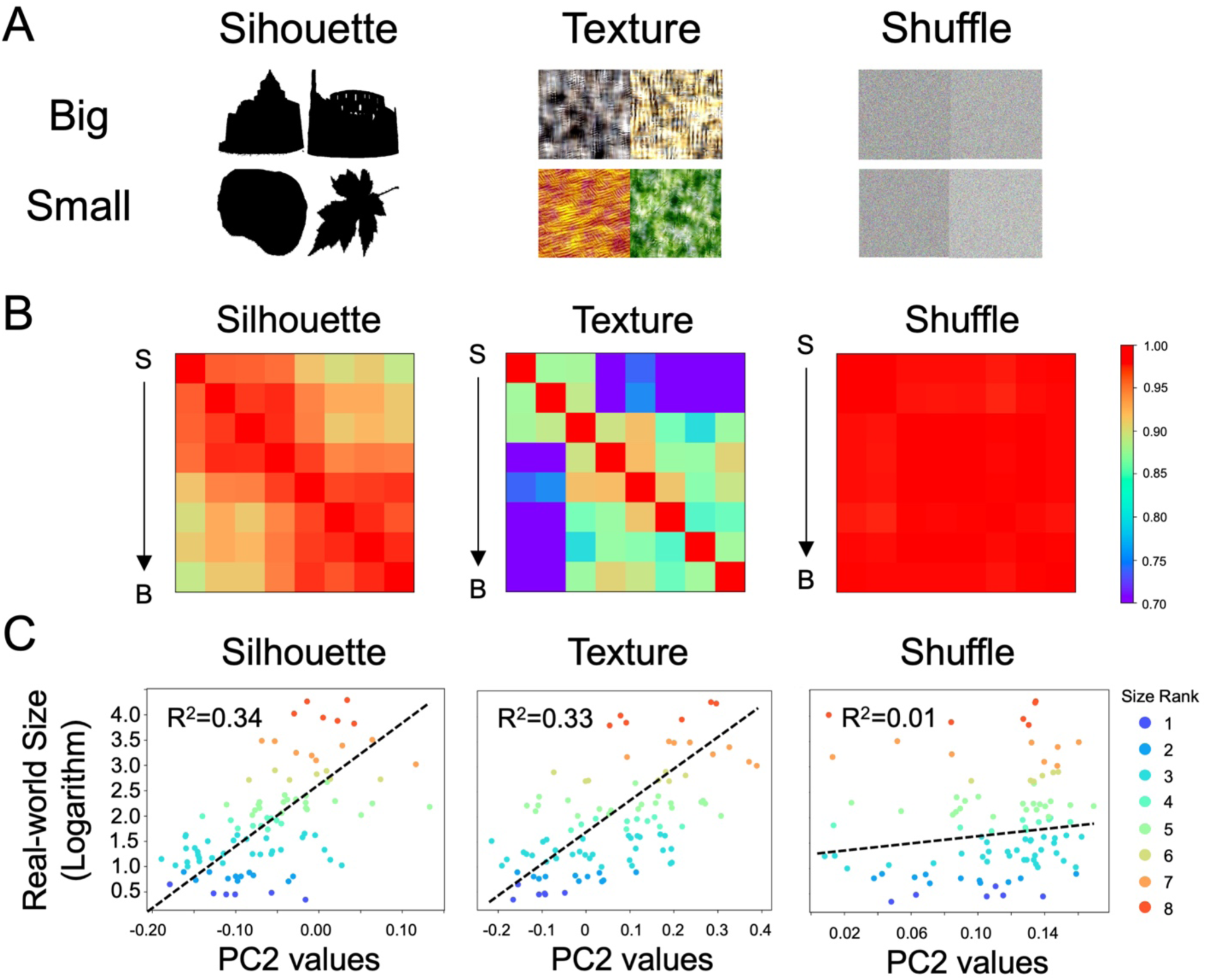
The intrinsic properties of objects’ shape and texture in representing the real-world size.(A) Examples of objects in silhouette, texture, and shuffle formats. (B) The RSM of Conv4’s responses to objects’ real-world size. The sensitivity to the real-world size was found when either the silhouettes or textures were presented, but not when the shuffle images were presented. (C) The natural logarithm correspondence between the PC2 and objects’ real-world size was found in the silhouettes and textures conditions, but not the shuffle condition. S: small objects; B: big objects.

An interesting question is whether humans used both shape and texture to represent objects’ real-world size in the brain. To address this question, we used fMRI to scan human subjects when they passively viewed the objects from the real-world size dataset and their silhouettes and textures. First, we replicated previous finding (*9*) with a medial-to-lateral arrangement of a big-to-small map separated by the middle fusiform sulcus (MFS) when the original objects were presented (Fig. 5A, top panel). Interestingly, only the silhouettes (Fig. 5A, middle panel), but not the textures (Fig. 5A, bottom panel), activated regions that are located along the medial-lateral transitions for distinguishing big or small objects. This observation was further quantified by a region-of-interest (ROI) analysis, where the ROIs were defined by the contrast of original big objects versus original small objects. We found that significantly higher BOLD responses to big objects were found in the silhouettes condition (t = 4.92, p < 0.001), but not in the texture condition (t = 1.14, p = 0.27) (Fig. 5B). Besides the univariate analysis, we also adopted a multivariate pattern analysis to examine whether the texture information was used to infer objects’ real-world size in humans. The multivariate pattern analysis in the ROIs revealed that only the BOLD responses to the silhouettes were able to distinguish big objects from small objects (classification accuracy = 0.80, p < 0.05; Fig. 5C). Finally, a whole-brain searchlight analysis failed to find any region that was capable of distinguishing the real-world size of objects based on the texture information (Fig. 5D). Therefore, although the DCNNs were able to use both shape and texture information to represent objects’ real-world size, in the human brain we only used the shape information.

**Fig. 5.**
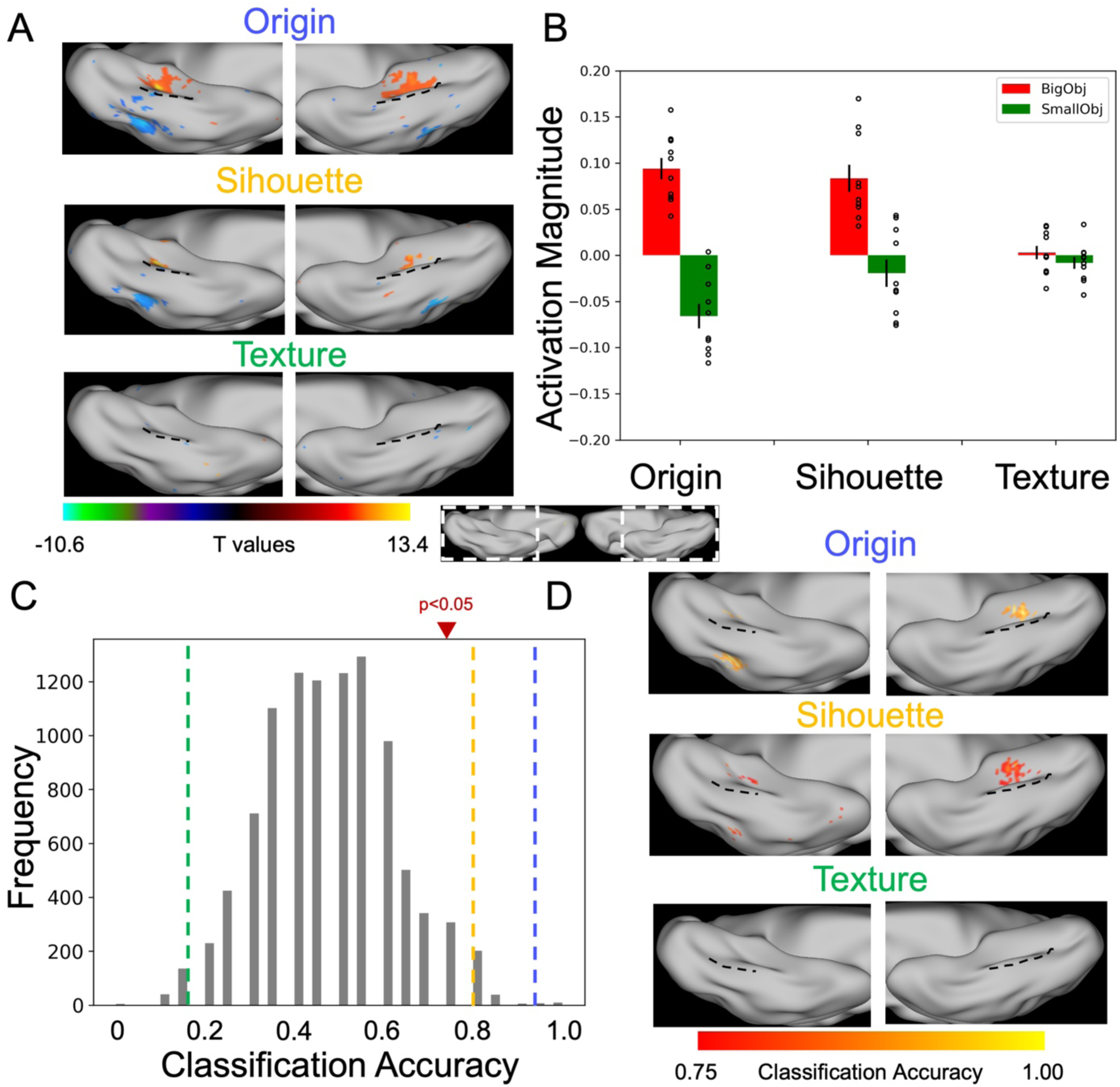
The properties of objects’ shape and texture in representing the real-world size in human brain. (A) Size-preference activation maps (big vs. small) from the group-averaged fMRI data when objects in original, silhouette, and texture formats. A medial-to-lateral arrangement of the real-world size preference in the ventral occipitotemporal cortex was present in the origin and silhouette conditions, but not in the texture condition. Black dotted line: middle fusiform sulcus (MFS). (B) Activation magnitude showed significant difference between big and small objects in the ROIs sensitive to objects’ real-word size in the silhouette condition, but not in the texture condition. (C) Multivariate pattern analysis found that neural response patterns in the origin and silhouette conditions were capable of distinguishing big objects from small objects, but not in the texture condition. Blue line: origin condition; Yellow line: silhouette condition; Green line: texture condition. (D) A whole-brain searchlight analysis found regions capable of distinguishing big from small objects only in the ventral occipitotemporal cortex and only for origin objects and silhouettes, but not for the texture condition.

The fMRI experiment on humans revealed that humans only used the shape information of objects to infer objects’ real-world size, whereas the DCNNs used both the shape and texture information of objects. The inconsistency between humans and DCNNs raised an interesting question of whether the texture information was necessary for the DCNN to use the feature of objects’ real-world size to construct object space. To address this question, here we used a stylized AlexNet (*23*), which was trained on a stylized version of ImageNet (i.e., Stylized-ImageNet). With randomly selected painting styles (Fig. 6A), the shape of objects was almost unchanged, while the texture was significantly distorted. We found that the stylized AlexNet showed a similar pattern as the original AlexNet, where Conv4’s responses showed sensitivity to objects’ real-world size (correlation with the ideal observer: r = 0.86, p < 0.001; Fig. 6B) and only the fourth axis of object space encoded this feature (Fig. 6C; DI = 0.14, p < 0.01, uncorrected) with the mapping function of the natural logarithm (R^2^ = 0.29, p < 0.001) (Fig. 6D).

**Fig. 6.**
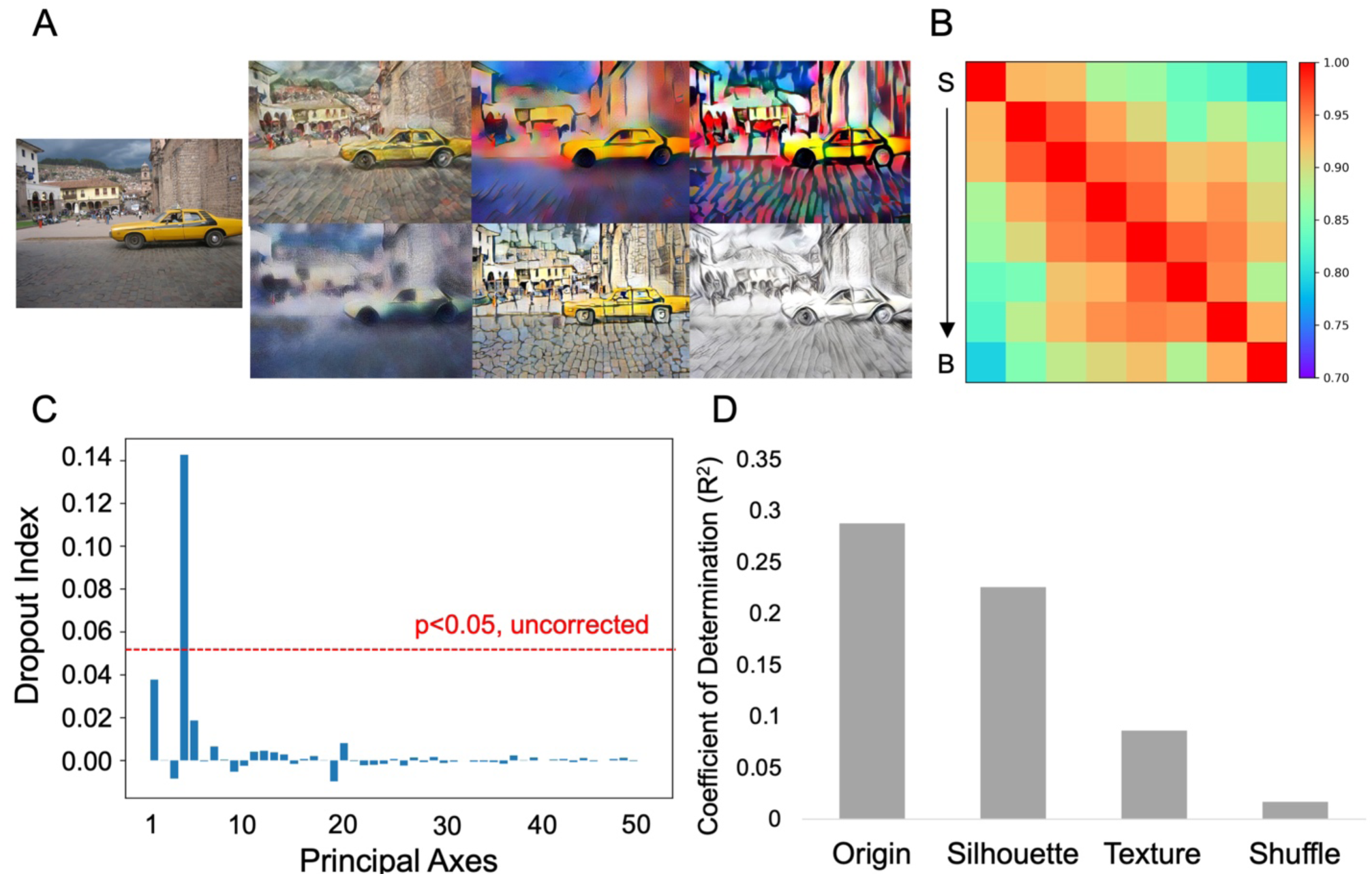
The necessity of texture information in DCNNs. (A) Images from the ImageNet were transformed with various painting styles, where the shape information was largely reserved and the texture information largely distorted. (B) The RSM of Conv4’s response to objects’ real-world size. (C) Only the fourth axis of object space constructed by the stylized AlexNet encoded the feature of objects’ real-world size. (D) Logarithm correspondence between the PC4 and objects’ real-world size showed less sensitivity to texture as compared to shape.

Importantly, this axis in object space was less sensitive to the texture information (R^2^ = 0.08, p < 0.001 for textures) as compared to the shape information (R^2^ = 0.22, p < 0.001 for silhouettes). In short, although the texture information was used in the DCNNs to infer objects’ real-world size, it was not necessary to construct an axis in object space to represent objects’ real-world size.

## Discussion

In this study, we used both DCNNs and fMRI to examine whether the real-world size of objects served as an axis of object space for object recognition. With the three criteria proposed here, we showed that to achieve successful performance in object recognition, the DCNNs automatically extracted information regarding the real-world size of objects (sensitivity criterion) to form an axis that was statistically independent of the rest axes of object space (independence criterion). Importantly, the removal of this axis significantly impaired DCNNs’ performance of object recognition (necessity criterion). Having identified the feature of objects’ real-world size as an axis of object space, we further performed *in silico* experiments to reveal that the intrinsic properties of the objects (i.e., shape and texture), rather than the external factors such as absolute retinal size, co-occurrence and task demands, were the key factors for DCNNs to infer the feature of objects’ real-world size. Finally, a comparative study between DCNNs and humans showed that humans only used the shape information, but not the texture information, to represent objects’ real-world size in the ventral occipitotemporal cortex of the brain. Taken together, by testing both DCNN and human subjects, our study provides the first comprehensive empirical evidence that the real-world size of objects served as an axis of object space, which illuminates the structure of object space in particular and the principle of processing objects in general.

The biological visual system needs to recognize a large number of objects with rich details within a very short time while retaining high accuracy in our daily life (*4, 24*). From the viewpoint of resource rationality (*25, 26*), it is unrealistic to process every detail when time is limited, and uneconomical because visual details are highly redundant and correlated. Object space is then proposed as a framework to simplify and abstract visual object representation by projecting critical features into a representational space with a finite number of orthogonal axes (*2, 4, 5, 27, 28*). The finite number of axes prevents the confusion of processing infinite visual details within a limited time, and statistical orthogonality among axes effectively reduces the redundancy of visual details (*29, 30*). Moreover, the existence of object space also permits perceptual invariance (*2, 28*) and generalization (*5*) during object recognition. Therefore, the identification of features encoded by axes of object space is uttermost to understand how objects are represented and recognized in the mind. Based on the framework of object space, we proposed that three criteria: sensitivity, independence and necessity, must be satisfied. However, limited by traditional methods, previous studies mainly focus on the first criterion of sensitivity that neuronal responses change as a function of feature magnitude, while leaving the rest of the two criteria unexamined. In this study, we used DCNNs as a model to examine the feature of objects’ real-world size as an axis of object space with all three criteria for the first time, which provides a novel approach for future exploration of other axes in object space.

Previous studies identifying features for object space mainly focus on the physical attributes of objects, such as physical appearance (e.g., having furs, wings), curvature (e.g., spiky, stubby), and conceptual knowledge (e.g., animate, artefacts). In contrast to the view of vision for perception, vision is also for action (*31*). In fact, objects’ real-world size not only describes the scale of an object in the natural environment (*7*), but also heuristically affects our action on objects (*32*). First, the space of the natural environment determines the layout of objects regularly. Smaller objects could be placed into larger objects, but not vice versa. This prerequisite influences our recognition of objects. For example, most cars are bigger than humans in real life, and when we see a car smaller than humans that violates the regularity of the object layout in the natural environment, we therefore need to recognize the car as a toy instead of a conveyance to accommodate this counterintuition. Second, the real-world size of objects provides heuristic information for affordance of how we manipulate objects. For example, small objects could be pinched, gripped or grasped, while larger objects could be held, lifted or raised. That is, the real-world size of objects intuitively guides our actions, with specific real-world size of an object associated with a set of specific actions, but not every action. Therefore, although a real car and a toy car share similar visual appearances, actions on these two objects are completely different (*8*). Further, the feature of objects’ real-world size also provides important information for high-level cognitive processes, such as attention (*33*), numerical perceptions (*34*), and semantic knowledge (*35*). In short, our finding that the feature of objects’ real-world size served as an axis of object space thus may greatly expand our understanding of object recognition from the angle of vision for perception to a novel viewpoint of vision for action.

Given the affordance property of objects, it is not surprising that external features on physical appearance such as co-occurrence of multiple objects and absolute retinal size, and features on levels of categorization (basic level versus superordinate level) had little effect on the representation of objects’ real-world size. Instead, the shape of objects, which is closely related to actions on objects (*31, 36, 37*), was found to be critical to construct the axis encoding objects’ real-world size in our study. A follow-up fMRI experiment confirmed that humans also used shape information to infer objects’ real-world size (*38–40*), suggesting that both artificial and biological systems adopted a similar algorithm to represent objects. One inconsistent finding, though, is that the DCNN also used texture information to infer objects’ real-world size whereas humans did not. In comparison to shape information, texture information reserves local features of objects, but not overall outlines. In real life, local features of objects were unstable in natural environments, largely affected by climates, air, and illumination. For example, the texture of a dog in the rain is dramatically different from the same dog rolling in the snow. Therefore, it is possible that human vision may consider the ecology of objects and therefore ignore the texture information of objects to infer objects’ real-world size. Consistent with the conjecture, by incorporating different painting styles, the texture of objects was largely unstable, and our stylized AlexNet trained with these images showed less sensitivity to the texture information of objects in constructing the axis for objects’ real-world size. Therefore, future DCNNs may consider making the models more ecologically feasible to behave more human alike.

## Materials and Methods

### Neural network models

Multiple pre-trained DCNNs were used in this study.

The AlexNet (*21*) includes 8 layers of computational units stacked into a hierarchical architecture; the first 5 layers are convolutional layers and the last 3 layers are fully connected layers. The second and fifth convolutional layers are followed by the overlapping max-pooling layers, while the third and fourth convolutional layers are connected directly to the next layer. Rectified linear unit (ReLU) nonlinearity was applied after all convolutional and fully connected layers. Layer 1 through 5 consisted of 64, 192, 384, 256, and 256 kernels.

Several extra DCNNs, including VGG11, VGG13, ResNet18, ResNet34, and Inception_v3, were used to verify whether results from the AlexNet could be replicated in other network architectures.

Two VGG networks (*41*), VGG11 and VGG13, were used to examine the effect of layer numbers on the formation of real-world size preference in DCNNs. The VGG11 and VGG13 include 11 and 13 layers respectively, with the first 8 and 10 layers being convolutional layers and the last 3 layers being fully connected layers. All hidden layers are equipped with the ReLU non-linearity. For VGG11, overlapping max-pooling layers follow the 1, 2, 4, 6, 8 convolutional layers. For VGG13, overlapping max-pooling layers follow the 2, 4, 6, 8, 10 convolutional layers.

Two ResNet networks (*42*), ResNet18 and ResNet34, were used to examine the effect of residue blocks on the formation of real-world size preference in DCNNs. ResNet18 and ResNet34 include 18 and 34 layers respectively, with all layers being convolutional layers except for the last one being a fully connected layer. A residue block was constructed between every two convolutional layers by inserting a shortcut connection. All hidden layers are also equipped with the ReLU non-linearity.

Inception_v3 (*43*) was used to examine the effect of inception structure on the formation of real-world size preference in DCNNs. Inception_v3 includes 5 independent convolutional layers, 10 Inception modules, and 1 fully-connected layer in total. Each Inception module consists of several convolutional layers with small kernel sizes arranged in parallel. The 10 Inception modules could be classified into InceptionA, InceptionB, InceptionC, InceptionD or InceptionE modules. Among the 10 Inception modules, three are InceptionA modules, followed by one InceptionB module, four InceptionC modules, one InceptionD module and two InceptionE modules. The detailed architecture could be referred to (*43*).

Each neural network model was pre-trained to perform object classification on the ILSVRC2012 ImageNet dataset (*44*), which includes about 1.2 million images of objects belonging to 1,000 categories. The object classification accuracy was evaluated on 50,000 validation images that were not seen by the model during training. The Top-1 and Top-5 accuracies of AlexNet are 52.6% and 75.1%. The network weights of all neural network models were downloaded from the PyTorch model Zoo (https://pytorch.org/vision/0.8/models.html) (*45*).

A stylized AlexNet (*23*), which has the same architecture as the classical AlexNet but is trained with a Stylized-ImageNet dataset (SIN) was downloaded from https://github.com/rgeirhos/texture-vs-shape/tree/master/models. The SIN was constructed by replacing the style (i.e., texture) of images from the ImageNet ILSVRC2012 training dataset with styles of different paintings (see Fig. 5A for examples). The object recognition ability of the stylized AlexNet was mainly achieved via shape but insensitive to texture variance of the object images.

### Stimulus datasets and model training

To examine whether real-world size preference emerges in the AlexNet pre-trained for object classification, we used a real-world size dataset downloaded from https://konklab.fas.harvard.edu/ImageSets/OBJECT100Database.zip, which was previously used (*22*) to evaluate real-world size preference in humans. This dataset includes 100 background-free object images spanning the range of real-world size from small objects (e.g., thumbtack) to large objects (e.g., colosseum). Each image consists of a single object, labeled with the measured real-world size of this object. The real-world size of each object was measured as the diagonal of its bounding box, ignoring the depth of the object, and quantified in centimeters.

Silhouette, texture, and shuffle versions of the real-world size dataset were separately generated to examine the contribution of shape and texture information to the representation of real-world size in AlexNet. In detail, to generate a silhouette of the original object, we first detect edges of an object using the canny edge detection algorithm, then set all values within edges as 0 (i.e., black color); this removed texture information from the original objects. On the other hand, a texture version of the original object was synthesized using the Portilla-Simoncelli algorithm (https://github.com/LabForComputationalVision/textureSynth) (*46*). In detail, the Portilla-Simoncelli algorithm obtained four sets of parameters from the target image and altered a random-noise image into a synthesized texture image by iteratively matching its parameter distributions with those of the target image. The four sets of parameters included 1) a series of first-order constraints on the pixel intensity distribution, 2) the local autocorrelation of the target image’s low-pass counterparts, 3) the measured correlation between neighboring filter magnitudes, and 4) cross-scale phase statistics. This procedure ensured that the random-noised images converged as a texture counterpart of the original images but reserved no shape information. The shuffle format of the real-world size dataset was generated by randomly shuffling the original images, which destroyed all shape and texture information of the object.

To evaluate whether the retinal size difference among objects accounted for real-world size preference of the AlexNet, we re-trained the AlexNet from scratch with a single-object version of the original ImageNet dataset that contains no background information. To do this, we first downloaded annotations of object bounding boxes from http://image-net.org/download-bboxes, which were annotated and verified through Amazon Mechanical Turk. 544,546 bounding boxes are corresponding to the original training images and 50,000 bounding boxes corresponding to the original validation images, respectively. We removed the background of each image by setting pixels outside the bounding box to 255 (i.e., white color). For images containing multiple bounding boxes, we randomly selected one bounding box as our target. The AlexNet was trained for 50 epochs, with an initial learning rate of 0.01 and a step multiple of 0.1 in every 15 epochs. Parameters of the model were optimized using stochastic gradient descent with the momentum and weight decay fixed at 0.9 and 0.0005. The Top-1 and Top-5 accuracies of the AlexNet that trained with the single-object image dataset (i.e., the single-object AlexNet) were 46.7% and 72.0%, respectively.

To evaluate the effect of task demand to the emergence of the real-world size axis in object space from the AlexNet, we re-trained an AlexNet from scratch to classify objects into two superordinate categories, the living things and artefacts (the AlexNet-Cate2). All object images were selected from the ImageNet dataset, which consisted of 866 categories in total. The number of images for training is 1,108,643, and for validating is 43,301. The AlexNet-Cate2 shared the same architecture as the original AlexNet, except that we added one extra FC layer after the FC3 layer for the classification of two superordinate categories. The AlexNet-Cate2 was trained following the same procedure as the single-object AlexNet. The Top-1 and Top-5 accuracies of the AlexNet-Cate2 were 94.7% and 100.0%, respectively.

### Calculate representational similarity matrix (RSM) of real-world size in AlexNet

Representational similarity between objects in different real-world sizes was used to evaluate whether real-world size preference automatically emerges in a DCNN that is trained for object recognition.

To achieve that, we extracted responses to objects from the real-world size dataset in different layers of the AlexNet. All images were transformed with resize and normalization to match the input requirement of the AlexNet. No ReLU was performed for the responses. We averaged responses from the convolutional layers within each channel, resulting in a response pattern of 256 channels for each image. We grouped objects into eight size ranks according to their real-world sizes (see Table S1). Each size rank included no less than six objects in different viewpoints, colors, and object shapes, to balance unrelated confounding factors. We averaged object response patterns within each size rank, and calculated similarity between these averaged response patterns in different size ranks to examine whether the response patterns in nearer size ranks is more similar with each other than those further apart, resulting in an RSM of size ranks in the AlexNet.

The degree of real-world size preference was quantified by comparing it with an ideal observer model. The RSM of the ideal observer model was constructed by the consistency between each pair of size ranks, which was defined as follows:

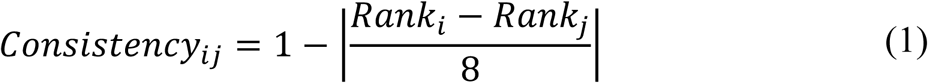

where *Rank*_*i*_ and *Rank*_*j*_ are size ranks, which ranged from 1 to 8. High correspondence between the RSMs of the AlexNet and the ideal observer suggested real-world size preference emerged in the AlexNet.

We also measured the real-world size representation of humans. Two participants (two males; 22 and 27 years) were recruited to separately compare the sizes of the objects from the real-world size dataset. For each pair of objects, participants were required to indicate which object is bigger than the other. Each participant completed 4,950 comparisons (i.e., 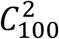) for all pairs of the 100 images, which provided a proportional value for each object with the following formula,

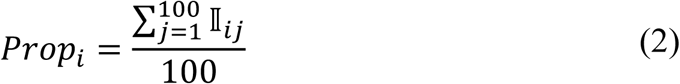

where 𝕀_*ij*_ equals to 1 when the *i*_*th*_ object is judged to be bigger than the *j*_*th*_ object. An object with higher proportional value indicated larger real-world size in a human’s mind. The proportional value of a size rank was calculated as the averaged proportional value across objects belonging to the same size rank. The RSM of humans was constructed by the consistency between each pair of size ranks measured as,

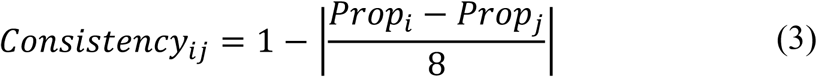

where *prop*_*i*_ and *prop*_*j*_ are proportional values of the *i*_*th*_ and *j*_*th*_ size rank.

### Evaluate the role of real-world size in object space

To evaluate whether real-world size played a role as a principal axis in object space, we used Principal Component Analysis (PCA) to recover an object space using images from the ImageNet validation dataset. Specifically, we first fed all 50,000 validation images into a pre-trained AlexNet. Responses from the layer with the highest real-world size preference (i.e., Conv4) were extracted and then averaged within each channel to generate a response matrix (Image numbers ×Channel numbers). We further normalized it by dividing its second-order norm across images. Then we used PCA to decompose the response matrix of the AlexNet into multiple principal axes. This yielded a linear transformation between responses and principal components (PCs) as follows,

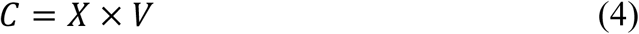

where C is the PCs, X is the responses and V is the principal axes in object space. We retained the first 50 principal axes, which captured 94.7% of the variance in the response of the AlexNet (More than 90% for other DCNNs).

We further investigated whether the real-world size was represented as a principal axis in object space. We first extracted responses to images from the real-world size dataset in the same layer that was used for the construction of principal axes, and got the projected principal components. To evaluate the effect of each axis on the real-world size preference, we iteratively removed the variance of each component from the original responses, and re-calculated the RSM of real-world size in the AlexNet. Reduction of correspondence to the ideal observer was measured with a dropout index (DI) defined as the following,

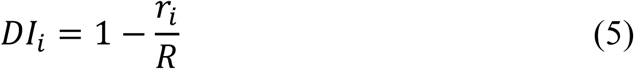

where *r*_*i*_ is the correspondence between the real-world size RSMs of AlexNet and the ideal observer after removing the *i*_*th*_ PC, and R is the correspondence without removing any PCs. Higher value indicated larger effect of a PC to the real-world size preference.

The significance of DIs was evaluated by comparing it with a null distribution, which was generated from an untrained AlexNet with repetition of the same procedures for 5,000 times for each principal axis. Multiple comparison correction was performed using Bonferroni correction after considering an integrated null distribution across all principal axes. The PC with significant DI value was identified as the axis encoding real-word size.

The quantitative relationship of this axis to the real-world size of objects was evaluated with a series of functions in different scales,

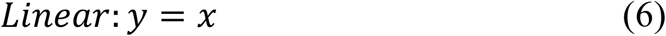

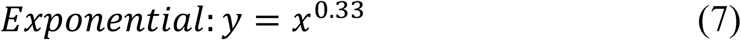

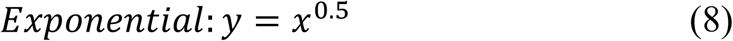

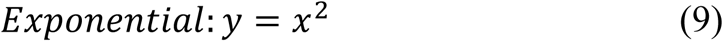

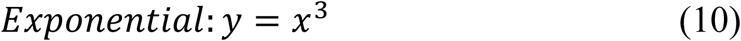

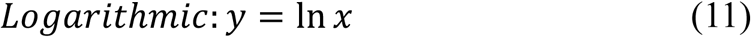

where x is the value of the real-world size PC, and y is the measured real-world size of objects. The scale with the best fit was taken as the relationship that best describes the data.

Necessity of the real-world size axis was evaluated with the performance of object recognition measured as the Top-1 categorization accuracy from the last fully connected layer. Similar to the measurement of DI, we first removed the variance of each PC from the original responses in Conv4, and then fed these responses into higher layers to get the Top-1 accuracy of the AlexNet for each category. Significance was also measured following the similar procedure as above. To reduce computational time, we iterated the procedure for 500 times to each category.

### fMRI Experiments

#### Participants

10 participants (5 males, age range: 18-27 years) from Beijing Normal University participated in this study to examine the effects of shape or texture on the real-world size representation in the human brain. All participants had a normal or corrected-to-normal vision. Informed consent was obtained according to procedures approved by the Institutional Review Board of Beijing Normal University.

#### Image Acquisition

Imaging data were collected on a 3T Prisma Siemens MRI Scanner with a 64-channel phased-array head coil at Beijing Normal University Imaging Center for Brain Research. The anatomical images were acquired with a magnetization-prepared rapid gradient-echo (MPRAGE) sequence. Parameters for the T1 image are: TR/TE=2,530/2.27ms, flip angle=7°, voxel resolution=1×1×1mm. Blood oxygenation level-dependent (BOLD) contrast was obtained with a gradient echo-planar T2* sequence. Parameters for the T2* image are: TR/TE=2,000/34.0ms, flip angle=90°, voxel resolution=2×2×2mm, FoV=200×200mm.

#### Experiment design

All participants completed eight runs of fMRI scanning. Participants were shown images of big or small objects in a standard block design. All objects were displayed at the same visual angle (5.3°×5.3°, visual distance=100cm) to exclude the confounding effect of different retinal sizes of objects. Big objects were selected as the largest 40 objects from the real-world size dataset, and small objects were selected as the smallest 40 objects from the same dataset. In addition, the same objects in a silhouette or texture were also used to evaluate the effect of shape and texture on the real-world size representation in the occipitotemporal cortex. Each run consisted of six conditions (i.e., big origin, small origin, big silhouette, small silhouette, big texture, and small texture). Each run lasted 320s, which included four block sets. Each block set lasted 60s, consisting of three conditions during which all 40 images were shown for each condition. Each image was presented for 200ms, followed by a 300ms fixation, consisting of a 500ms trial. The position of each object on the screen was slightly jittered to increase the attention of the participant. The first two block sets presented images from all of the six conditions without repetition, and the last two block sets presented images in palindrome. Five fixation periods of 16s intervened between each block set. Participants were instructed to pay attention to the images and to press a button when a red frame appeared around an object, which appeared twice per condition randomly.

#### Preprocessing

Anatomical and functional data were preprocessed using fMRIPrep (version 20.2.0) (*47*). Pre-processing included skull stripping, slice-time correction, co-registration to T1w with boundary-based registration cost-function, correction for head-motion and susceptibility distortion, and temporal high-pass filtering (128s cut-off). All structural and functional images were projected into a 32k_fs_LR space (*48*) using the ciftify toolbox (*49*). The functional images were spatially smoothed with a 4 mm FWHM kernel.

#### Univariate Analyses

First-level statistical analyses were performed for functional images of each participant in each run using the general linear model (GLM) from the HCP Pipelines. Second-level statistical analyses were separately performed on odd or even runs of the activation maps generated from the first-level analysis.

Activation magnitude was measured as the Cohen’s d with the following formula,

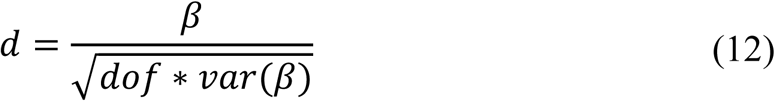

where *β* is the contrast of parameter estimation (COPE), and *dof* is the degree of freedom.

Whole-brain group analysis across participants was conducted based on the second level statistical maps of even runs using the Permutation Analysis of Linear Models (PALM) (*50*). A contrast was performed at an uncorrected threshold of p < 0.005 to identify the regions selectively active to original big objects versus original small objects. The identified regions were used as a mask of regions sensitive to real-world size in the following analyses.

#### Multivariate analyses

Multivariate analyses used data from odd runs. We assessed whether the multivoxel patterns of different conditions (i.e., origin, silhouette, and texture) in the mask of the real-world size were sufficient to classify the size category (i.e., big or small) of the object being viewed. The classification was performed on a support vector machine (SVM) with a linear kernel using the leave-one-out cross-validation (LOOCV) across participants, and its accuracy was evaluated as the averaged accuracies from all runs of the LOOCV. The significance of classification accuracy was evaluated with a null distribution, which was built by classifications after permutating the pooled activations from the original conditions 10,000 times. Significance (p < 0.05) was achieved when classification accuracy was larger than 0.75. Searchlight analysis was also performed to test for coding of real-world size in the whole brain. The same procedure of classification was evaluated in small spherical ROIs (radius 10 mm) centered on each vertex of the brain in turn. This generated three whole-brain maps corresponding to original objects, silhouettes, and textures. The searchlight maps were spatially smoothed using a Gaussian smoothing kernel with FWHM as 2mm.

## Author contributions

T.H., Y.S and J.L. designed research and wrote the manuscript. T.H. performed research and analyzed data. All authors contributed to the article and approved the submitted version.

## Data and materials availability

All code and data underlying our study and necessary to reproduce our results are available on Github: https://github.com/helloTC/RealWorldSizeAxis. The datasets presented in this study can be found in online repositories, the names of each repository and the download location can be found in the article.

## Supplementary Materials

**Fig. S1.**
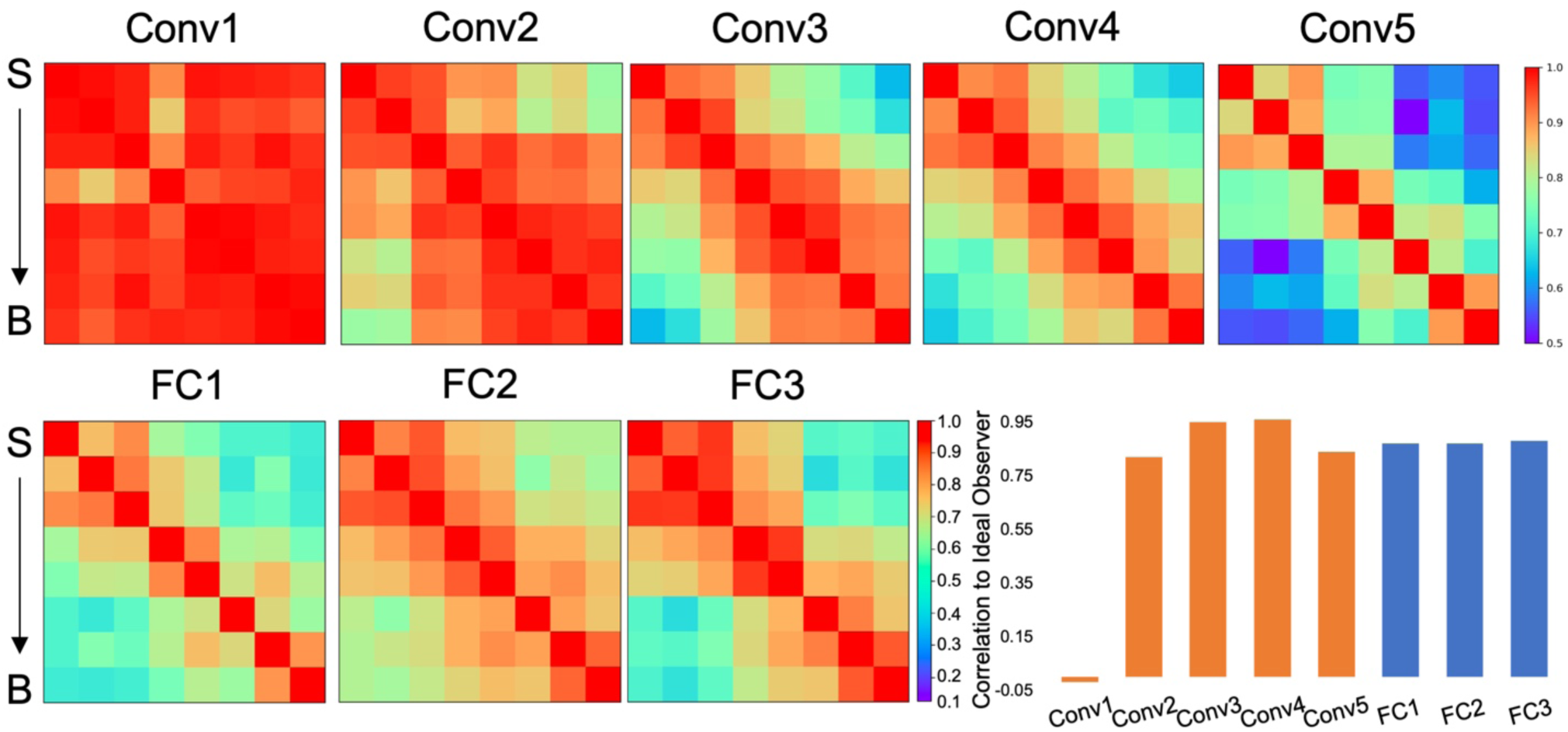
Representational similarity matrix (RSM) of size ranks in different layers of AlexNet. The RSM showed high correspondence to the ideal observer from the Conv2 layer, and achieved the highest value in the Conv4 layer. S: small objects; B: big objects.

**Fig. S2.**
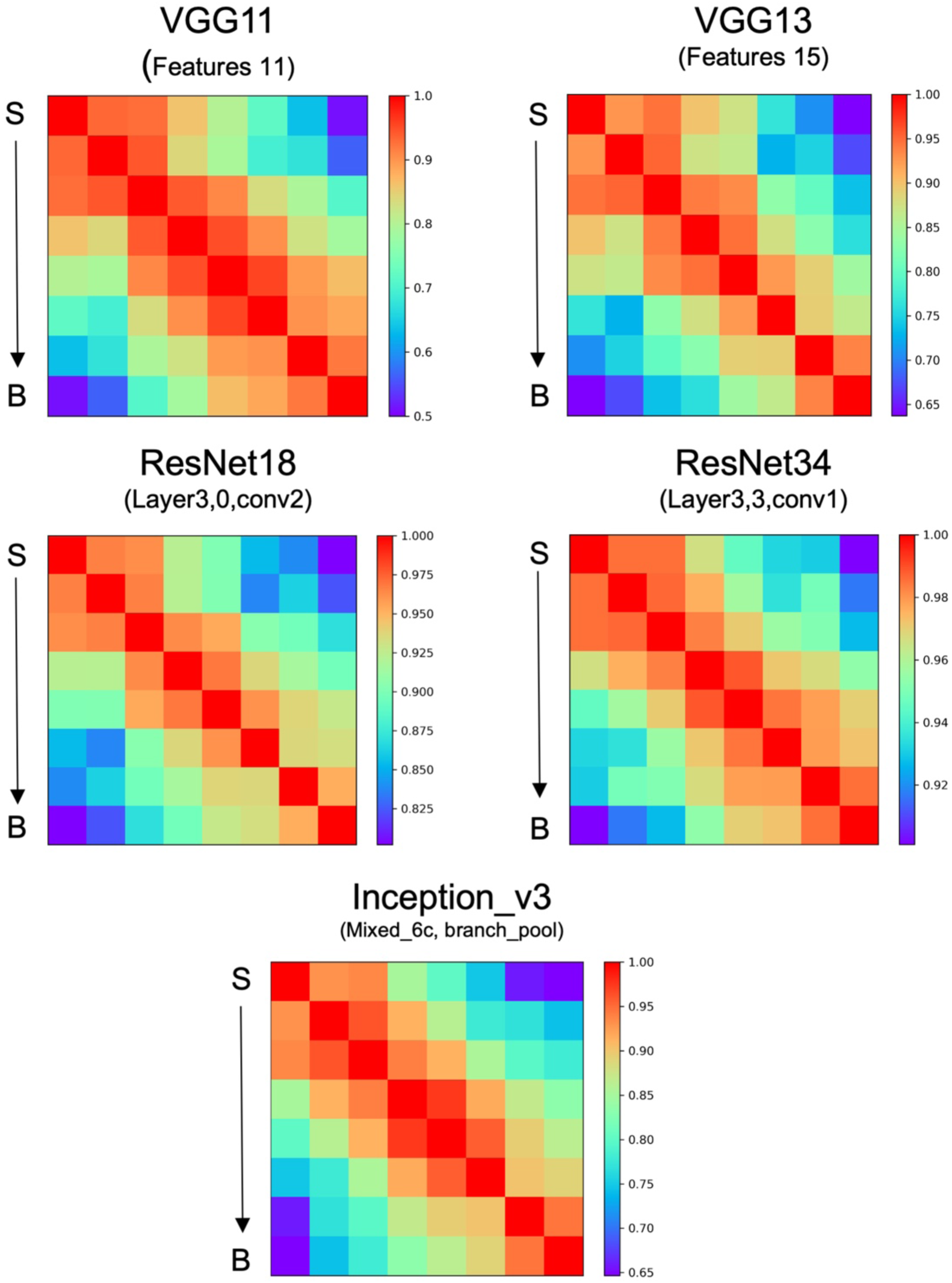
Preference to objects’ real-world size could be consistently found in DCNNs with different architectures. Five representative implementations, including two from the VGG family, two from the ResNet family, and one from the Inception family were examined. Their correlations to the ideal observer are 0.95 (VGG11), 0.96 (VGG13), 0.96 (ResNet18), 0.96 (ResNet34) and 0.97 (Inception_v3), respectively. S: small objects; B: big objects.

**Fig. S3.**
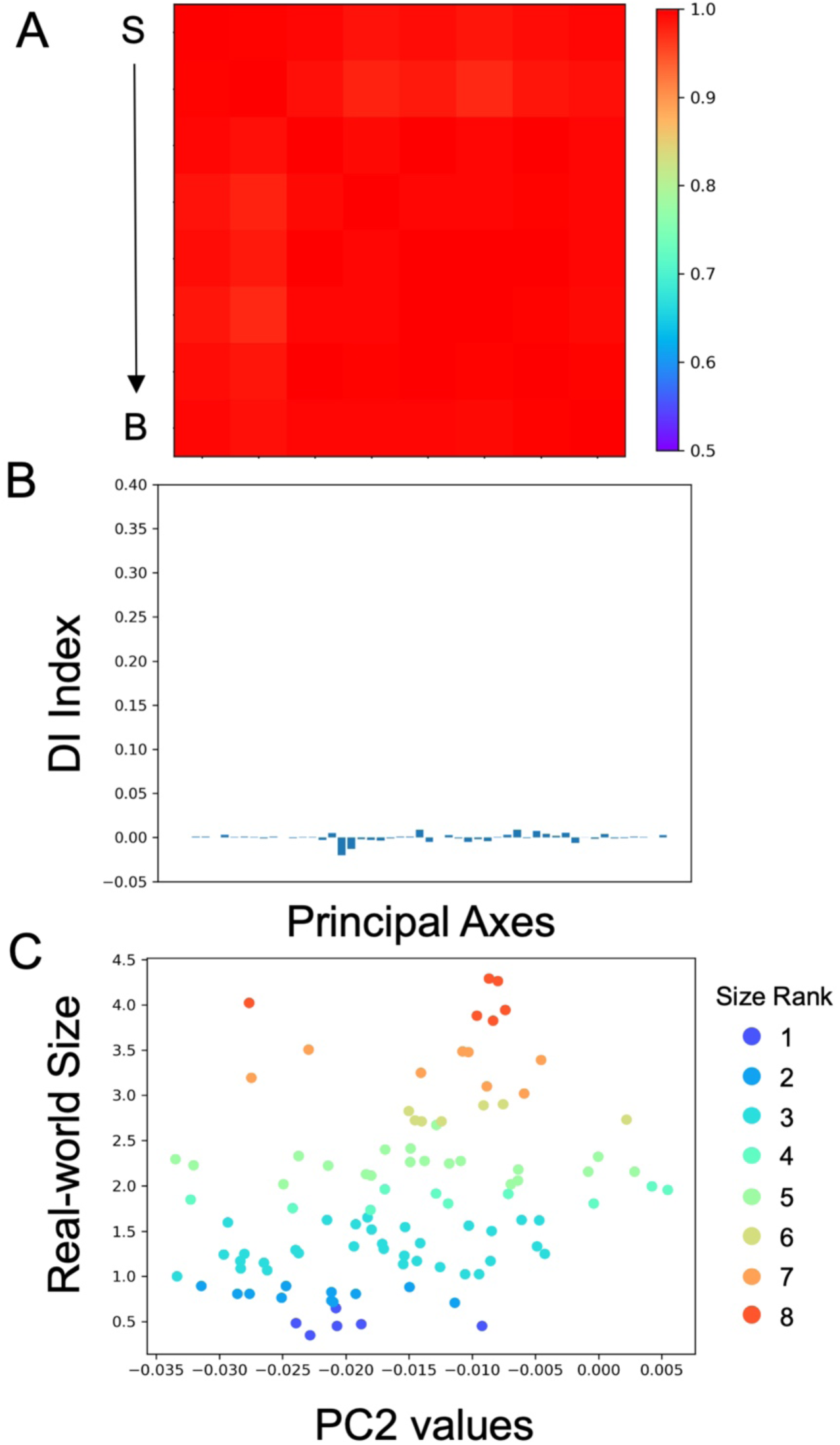
Untrained AlexNet showed no preference for objects’ real-world size. (A) The RSM of size ranks in untrained AlexNet could not distinguish object sizes. (B) No principal axis encoded the real-world size of objects in untrained AlexNet. (C) No logarithm correspondence between the PC2 values and the real-world size of objects. S: small objects; B: big objects.

**Fig. S4.**
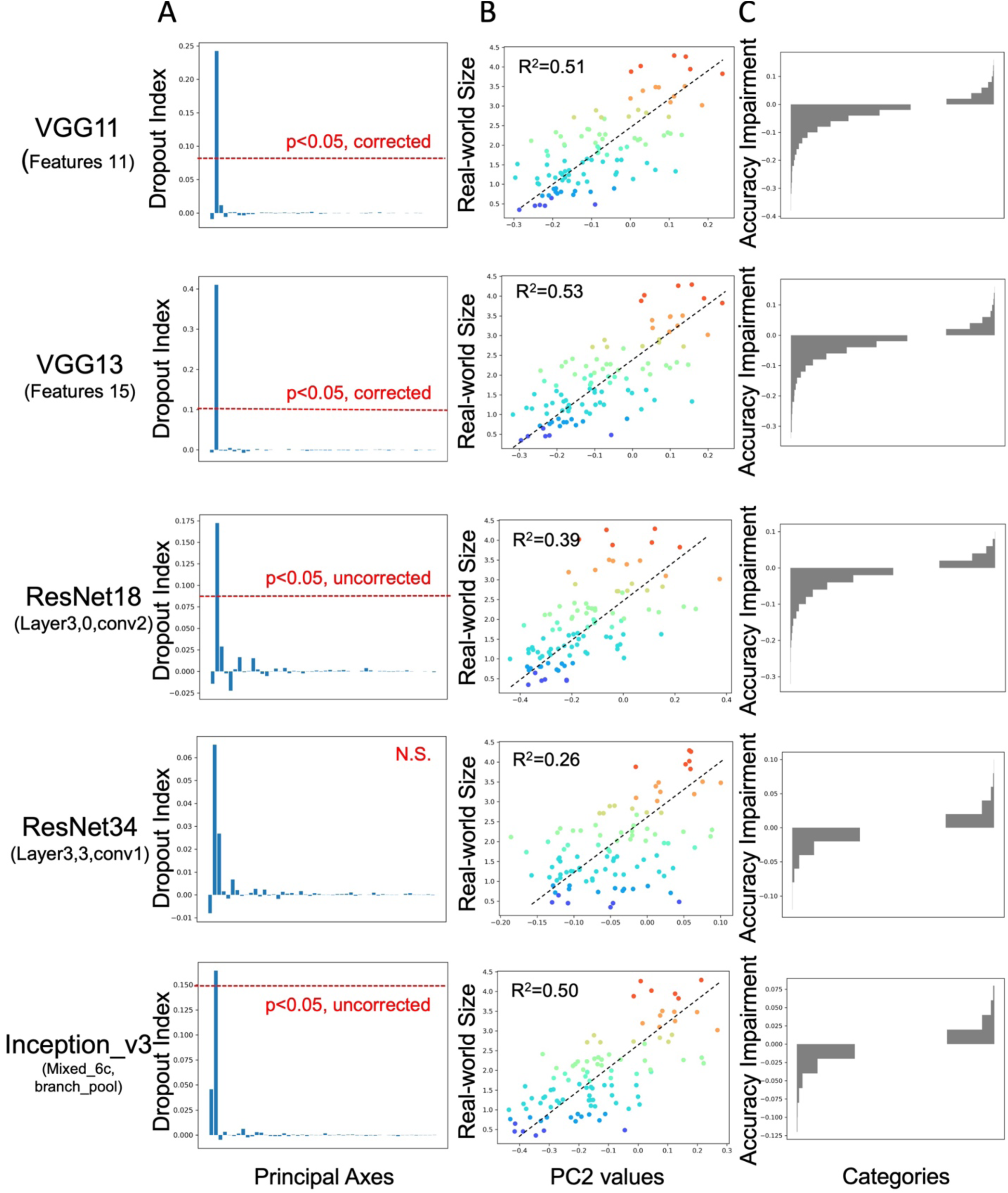
The real-world size axis emerged in object space in DCNNs with different architectures.(A) The size axis could be found in five different DCNNs, including two VGG networks, two ResNet networks, and one Inception network. The size axis from the ResNet34 was not significant while had a tendency. (B) Logarithm correspondence between the PC2 and real-world size of objects of the five DCNNs. (C) Impairment of prediction accuracy after removing variance that aligned to the size axis.

**Fig. S5.**
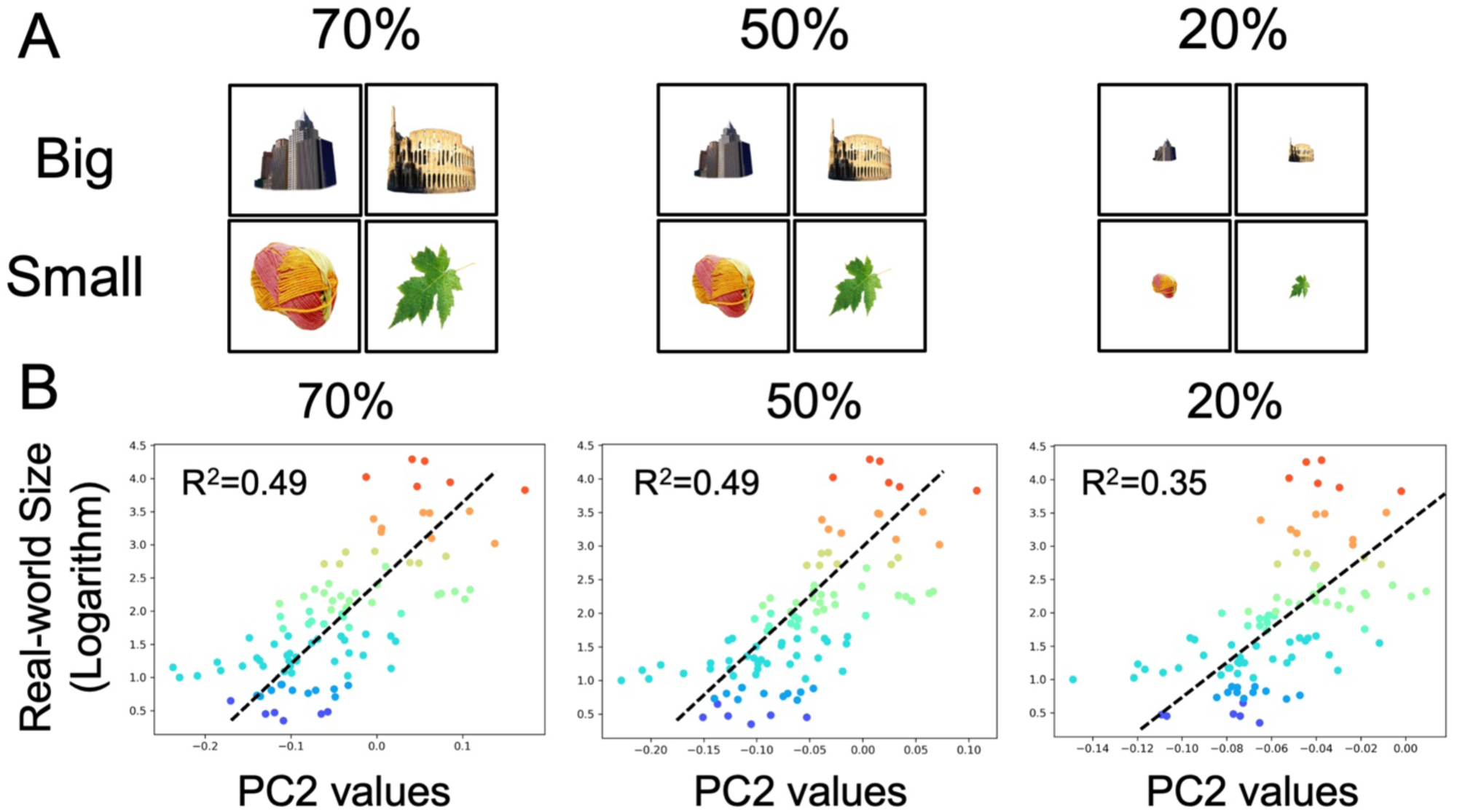
The size axis in object space has tolerance to the retinal size of objects. (A) Illustration of stimuli used for testing whether retinal size accounted for the real-world size axis in object space. Objects were narrowed as the 70%, 50%, 20% of the original images. (B) Natural logarithm correspondence between the PC2 and real-world size was relatively stable.

**Table S1.** Size ranks of objects. The 100 background-free objects were grouped into eight size ranks to control confounding factors which was unrelated to real-world size. All size ranks contain no less than six objects to guarantee unrelated factors could be removed as much as possible. The unit of size range is centimeter.

| Size Rank | Size Range | Number of Objects | Representative objects |
| --- | --- | --- | --- |
| 1 | 1-5 | 6 | Button, Dice, Thumbtack |
| 2 | 5-10 | 11 | Key, Garlic, Teabag |
| 3 | 10-50 | 32 | Shoe, Mouse, Helmet |
| 4 | 50-100 | 10 | Backpack, Screen, Cooler |
| 5 | 100-500 | 20 | Grill, Refrigerator, Sofa |
| 6 | 500-1,000 | 7 | Car, Statue, Fountain |
| 7 | 1,000-5,000 | 8 | Monument, Lighthouse, Freight train |
| 8 | 5,000+ | 6 | Airplane, Colosseum, Arch of Triumph |

